# *RNA-Seq-Pop:* Exploiting the sequence in RNA-Seq - a Snakemake workflow reveals patterns of insecticide resistance in the malaria vector *Anopheles gambiae*

**DOI:** 10.1101/2022.06.17.493894

**Authors:** Sanjay C Nagi, Ambrose Oruni, David Weetman, Martin J Donnelly

## Abstract

**Background:** We provide a reproducible and scalable Snakemake workflow, called *RNA-Seq-Pop,* which provides end-to-end analysis of RNA-Seq data sets. The workflow allows the user to perform quality control, differential expression analyses, call genomic variants and generate a range of summary statistics. Additional options include the calculation of allele frequencies of variants of interest, summaries of genetic variation and population structure (in measures such as nucleotide diversity, Watterson’s θ, and PCA), and genome wide selection scans (*F_st_*, PBS), together with clear visualisations. We demonstrate the utility of the workflow by investigating pyrethroid-resistance in selected strains of the major malaria mosquito, *Anopheles gambiae.* The workflow provides additional modules specifically for *An. gambiae,* including estimating recent ancestry and determining the karyotype of common chromosomal inversions.

**Results:** The Busia lab-colony used for selections was collected in Busia, Uganda, in November 2018. We performed a comparative analysis of three groups: a parental G24 Busia strain; its deltamethrin-selected G28 offspring; and the susceptible reference strain Kisumu. Measures of genetic diversity reveal patterns consistent with that of laboratory colonisation and selection, with the parental Busia strain exhibiting the highest nucleotide diversity of 1.04·10^-3^, followed by the selected Busia offspring (7.1·10^-4^), and finally, Kisumu (6.2·10^-4^). Differential expression and variant analyses reveal that the selected Busia colony exhibits a number of distinct mechanisms of pyrethroid resistance, including the *Vgsc-995S* target-site mutation, upregulation of SAP genes, P450s, and a cluster of carboxylesterases. During deltamethrin selections, the 2La chromosomal inversion rose in frequency (from 33% to 86%), suggesting a link with pyrethroid resistance, which was previously observed in field samples from the same region. *RNA-Seq-Pop* analysis also reveals that the most widely-used insecticide-susceptible *An. gambiae* strain, Kisumu, appears to be a hybrid strain of *An. gambiae* and its sibling species *An. coluzzii*, which should be taken into consideration in future research.

*RNA-Seq-Pop* is designed for ease of use, does not require programming skills and integrates the package manager Conda to ensure that all dependencies are automatically installed for the user. We anticipate that the workflow will provide a useful tool to facilitate reproducible, transcriptomic studies in *An. gambiae* and other taxa.

## Introduction

Transcriptomics is central to our understanding of how genetic variation influences phenotype (Stark et al., 2019). In recent years, RNA-Sequencing has replaced microarray technologies for whole-transcriptome profiling, providing a relatively unbiased view of transcript expression (Zhao et al., 2014) with associated higher sensitivity and greater dynamic range (Lowe et al., 2017). The utility of RNA-seq is exemplified by the vast amounts of data accruing (Van den Berge et al., 2019), and in the many discoveries it has revealed – such as the extent of alternative splicing, and the biology of non-coding RNAs (Stark et al., 2019; Wang et al., 2010; Wang & Burge, 2008).

In recent years, various computational workflows have been developed to analyse RNA-Seq data in a reproducible manner (Lataretu & Hölzer, 2020; Zhang & Jonassen, 2019), however, these workflows are designed with the primary aim of differential expression analysis (DEA) and leave a large amount of untapped sequence-based information. In our own area of research, vector genomics, a scan of the literature revealed thirty-three RNA-Sequencing studies (supplementary table 1), of which only five interrogated the sequence data (Bonizzoni et al., 2015; David et al., 2014; Faucon et al., 2017; Kang et al., 2021; Messenger et al., 2021). A barrier to exploiting the full range of information contained within RNA-Seq data sets has been the absence of comprehensive, user-friendly pipelines which permit easily reproducible analysis (Grüning et al., 2018) and enable comparisons across studies.

In this study, using the workflow management system Snakemake (Mölder et al., 2021), we present a reproducible computational workflow, *RNA-Seq-Pop,* for the analysis of Illumina RNA-Sequencing datasets. The workflow is applicable to any paired-end Illumina RNA-Sequencing data. However, we also present modules specifically of interest in the analysis of the major malaria mosquito, *Anopheles gambiae s.l.,* and demonstrate their use in a study of pyrethroid-resistance in a strain of *An. gambiae* from Busia, Uganda.

Pyrethroids are the most widely used class of insecticide in malaria control, and over the past two decades, resistance in malaria vectors has spread throughout sub-Saharan Africa, posing a threat to vector control efforts (Ranson, 2017). In this period, the incrimination of genes involved in insecticide-resistant phenotypes of *Anopheles gambiae* has been primarily based on transcriptomic studies. For many years, these were performed using microarrays; synthesis of which has highlighted the repeatable overexpression of a handful of genes involved in detoxification, confirming well-established cytochrome P450s as candidates, whilst also implicating more diverse genes such as ABC transporters and sensory appendage proteins (Ingham et al., 2018). Yet to date, relatively few diagnostic markers have been identified, and important genes have been missed by standard transcriptomic analyses (Njoroge et al., 2021). These shortcomings illustrate the need for a more comprehensive approach to marker discovery. While wholegenome sequencing is providing valuable information on known and novel resistance variants (Clarkson et al., 2021; The *Anopheles gambiae* 1000 Genomes Consortium, 2020) exploiting the sequence data within RNA-Seq can help bridge the step from transcriptomics to genomics.

In Uganda, pyrethroid resistance has escalated in recent years (Lynd et al., 2019; Tchouakui et al., 2021). As well as the *Vgsc-995S* mutation, which has repeatedly been associated with pyrethroid-resistance, recent genomic studies from this region have shown that a triple-mutant haplotype, linking a transposable element, a gene duplication (*Cyp6aa1)* and a non-synonymous mutation *Cyp6p4-I236M,* is an important marker of pyrethroid resistance (Njoroge et al., 2021). A SNP-array based GWAS also demonstrated the *Cyp4J5-L43F* mutation to be a useful marker for insecticide resistance, whilst also implicating the 2La inversion karyotype as a potential marker (Weetman et al., 2018). We use *RNA-Seq-Pop* to uncover patterns of insecticide resistance in Ugandan *An. gambiae,* monitoring these resistance-associated mutations, whilst performing differential expression analyses, summarising genetic variation and ancestry, and karyotyping chromosomal inversions.

## Materials & Methods

### *RNA-Seq-Pop* implementation

We designed the *RNA-Seq-Pop* workflow according to Snakemake best practices (Köster, 2022). *RNA-Seq-Pop* is constructed with a single configuration file in human-readable yaml format (the config file), alongside a simple tab-separated text file containing sample metadata (the sample sheet). The overall *RNA-Seq-Pop* workflow is shown in figure 1.

**Figure 1:**
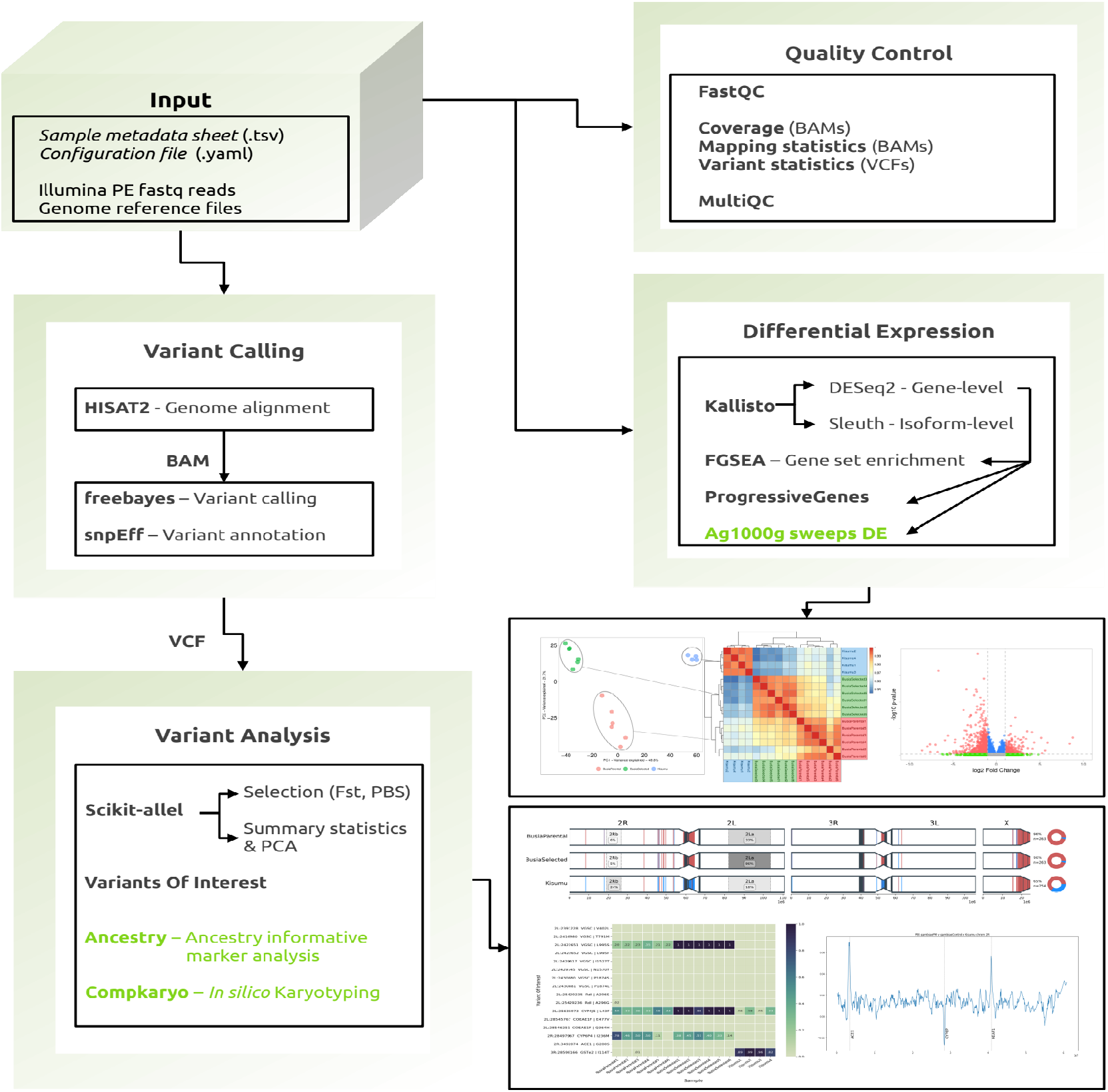
The *RNA-Seq-Pop* workflow and example outputs. The workflow has been designed for ease of use, requiring only a configuration file to set up workflow choices and a sample sheet to provide sample metadata. Modules highlighted in green are specific to *An. gambiae s.l.*

Dependencies are internally managed by the package manager Conda; to install all required software, specify the --use-conda directive at the command line, and Conda will automatically create isolated software environments in which to run. As of v1.0.0, *RNA-Seq-Pop* modules are written in Python (75% of the codebase) and R (25%), and internally, the workflow utilises a library (RNASeqPopTools) which providethe infrastructure to the Python codebase, to ensure readability. We provide a tutorial in the GitHub wiki to guide users on how to set up and run *RNA-Seq-Pop*.

#### Quality Control

The workflow begins by checking concordance between the user-provided sample metadata, configuration file, and reference and fastq files. Quality control metrics of fastq files are calculated with fastqc (Andrews, 2010), and logs and statistics from eight tools in the workflow are integrated into a report with MultiQC (Ewels et al., 2016). Raw fastq reads may be optionally trimmed with cutadapt (Martin, 2011).

#### Differential expression

Trimmed reads are aligned to the reference transcriptome with *Kallisto* (Bray et al., 2015) and differential expression performed at the gene-level with *DESeq2* (Love et al., 2014) and at the isoform-level with *sleuth*(Pimentel et al., 2017). The gene-level counts are normalised to account for sequencing depth, and principal components analysis (PCA) and Pearson’s correlation performed among all samples, and on subsets of the user-selected treatment groups used in differential expression analysis. Plots of these analyses are useful for exploratory data visualisation, providing an additional quality control step to ensure expected relationships between samples. *RNA-Seq-Pop* combines differential expression results from multiple pairwise comparisons into a Microsoft Excel spreadsheet for the user, as well as generating individual .csv files, volcano plots, and identifying the number of differentially expressed genes at various FDR-adjusted p-value thresholds. We use the R package *fgsea* (Korotkevich et al., 2021) for GO term and KEGG pathway enrichment analysis, on the most highly ranked genes based on differential expression (FDR-adjusted) p-values and, and optionally *F_ST_* values.

#### Variant calling

Reads are aligned to the reference genome with *HISAT2* (Kim et al., 2019) and read duplicates marked with *samblaster* (Faust & Hall, 2014) producing binary alignment files (BAM), which are sorted by genomic coordinate and indexed with *SAMtools* v1.19 (Danecek et al., 2021). SNPs are then called with the Bayesian haplotype-based caller *freebayes* v1.3.2 (Garrison & Marth, 2012). SNPs are called jointly on all samples, with different treatment groups called as separate populations, at the ploidy level provided by the user in the configuration file. The workflow internally parallelises *freebayes* by splitting the genome into small regions, greatly reducing overall computation time. The separated genomic regions are then concatenated with *bcftools* v1.19 (Danecek et al., 2021) and the final VCF piped through *vcfuniq* (Garrison et al., 2021), to filter out any duplicate calls that may occur at the genomic intervals between chunks. Called variants are then annotated using *snpEff* v5.0 (Cingolani et al., 2012).

#### Variant analysis & selection

*RNA-Seq-Pop* can then perform analyses on the variants called by *freebayes.* We apply filters to the data, including restricting to SNPs (excluding indel calls) and applying missingness and quality filters. We recommend using a quality score of 30 and a missingness proportion of 1, meaning a variant call (reference or alternate allele) must be present in each sample, i.e there are no missing allele calls. For each pairwise comparison specified in the config file, the workflow can perform a windowed Hudson’s *F_ST_* scan (Bhatia et al., 2013; Hudson et al., 1992) along each chromosomal arm, outputting windowed *F_ST_* estimates and genome-wide plots. Population branch statistic (PBS) scans may also be performed, conditional on the presence of three suitable populations for the phenotype(s) of interest (Yi et al., 2010). It is also possible to run Hudson’s *F_ST_* and PBS scans, taking the average for each protein-coding gene, rather than in windows. All population genetic statistics are calculated in scikit-allel v1.2.1. (Miles & Harding, 2017). We also provide a script (geneScan.py) to interrogate the VCF files, reporting missense variants from any gene of the user’s choice. A tab-separated file of variants of interest can be provided, from which the workflow will produce allele frequency heatmaps for each biological replicate and averaged across treatment groups. We define the expressed allele balance as the allele frequency at a genomic location in the aligned read data – for this analysis, *RNA-Seq-Pop* does not use variants called by *freebayes,* but instead calculates the proportion of each allele directly in bam files. An example variant of interest file for *An. gambiae* is provided in the *RNA-Seq-Pop* GitHub repository.

All analyses described thus far can be conducted across all taxa of any ploidy, requiring only a reference genome (.fa), transcriptome (.fa), and genome annotation files (.gff3).

#### Anopheles gambiae s.l specific analyses

For *Anopheles gambiae s.l* datasets we have exploited the *Anopheles gambiae* 1000 genomes resource (Miles et al., 2017; The *Anopheles gambiae* 1000 Genomes Consortium, 2020), to incorporate H12 and iHS (Garud et al., 2015) genomic selective sweep analysis. The workflow outputs the differentially expressed gene’s genomic location, the specific sweep signals present in the Ag1000g resource at that genomic location, and whether the region is a known insecticide resistance-associated locus. We caution that this kind of analysis is exploratory: many genes will be contained within selective sweeps, and may not have a causal link to phenotypic variation.

#### Population structure, ancestry and karyotyping

To investigate population structure, we apply SNP quality and missingness filters to the SNP data, which can be configured by the user. Multiple measures of population genetic diversity are estimated for each sample, such as nucleotide diversity (π), Watterson’s θ (Watterson, 1975), and inbreeding coefficients. We then prune SNPs in high linkage by excluding variants above an R^2^ threshold of 0.01 in sliding windows of 500 SNPs with a step size of 250 SNPs, and perform a PCA on the remaining SNPs. If the analysed species is *An. gambiae, An. coluzzii,* or *An. arabiensis,* the pipeline can implement an analysis of putative ancestry informative markers (AIMs). The AIMs were derived from two different datasets. The *An. gambiae/An. coluzzii* AIMs derive from the 16 genomes project (Neafsey et al., 2015) and in West Africa may distinguish between individuals with *An. gambiae* or *An. coluzzii* ancestry. The *An. gambcolu/An. arabiensis* AIMs are derived from phase 3 of the *Anopheles gambiae* 1000 genomes project, and distinguish between individuals with either *An. gambiae* or *An. coluzzi* ancestry from *An. arabiensis.* The relative proportion of ancestry is reported and visualised for the whole genome by chromosome. We modified the program *compkaryo* (Love et al., 2019) to enable the identification of common inversions on chromosome 2 in pooled samples.

### Busia RNA-Seq

#### Mosquito lines

We used a pyrethroid-resistant colony of *Anopheles gambiae s.s* from Busia, Uganda, alongside the standard multi-insecticide-susceptible reference strain, Kisumu. After 24 generations in colony, we stored RNA from the Busia strain (Busia parental), and selected the remaining colony using 0.05% deltamethrin papers in WHO tube assays for 4 generations (full details of the selection regime can be found in the supplementary text 2). We exposed females from the selected generation (G28) for one hour to 0.05% deltamethrin WHO papers using standard protocols, left for 24 hours post-exposure, and survivors were stored at −80°C prior to RNA extraction (Busia selected). Unexposed, age-matched Kisumu females were used as controls and stored in −80°C prior to RNA extraction.

#### Library prep

We extracted RNA from pools of five, 4-day old female mosquitoes using a Picopure RNA isolation kit (Arcturus, Applied Biosystems, USA). We performed six replicates for each Busia-derived treatment group, and four for Kisumu. Library quality and quantity were determined on a Tapestation 2200 (Agilent, UK) using high sensitivity RNA screentape. Paired-end 150bp RNA-Sequencing libraries were prepared and sequenced by Novogene (https://en.novogene.com/), on an Illumina NovaSeq 6000 system.

## Results

### Busia resistance phenotyping

The parental G24 *An. gambiae* Busia strain had lost much of its pyrethroid resistance during the time in culture and exhibited susceptibility to deltamethrin (100% mortality, 96.3-100 95% CIs) and low-prevalence resistance to permethrin (92.6% mortality, 85.6-96.4 95% CIs). Four generations of deltamethrin selections, demonstrated this loss to be readily reversible and resulted in a G28 selected Busia strain that showed increased resistance to both deltamethrin (69.7% mortality, 63.2-75.6 95% CIs) and permethrin (21.7% mortality, 14.9-30.5 95% CIs) when exposed for one hour in WHO tube assays. We compared the two Busia strains to one another, and to the pyrethroid-susceptible reference strain, Kisumu.

### RNA-Sequencing

As an illustrative example of the modules and output of the *RNA-Seq-Pop* workflow, we will describe the analysis of the Busia RNA-Seq dataset.

#### Quality control

We used *RNA-Seq-Pop* to import FASTQ data files into FastQC (Andrews, 2010) to determine levels of adaptor content, quality scores, sequence duplication levels and GC content in the raw read data. After genome alignment, we applied rseqQC and SAMtools to collect mapping statistics from the resulting BAM files. We then integrated MultiQC into the workflow, which collates statistics and results from eight tools to generate a convenient, interactive (.html) quality control report. Figure 2 shows reports generated by multiQC on the Busia *An. gambiae* dataset.

**Figure 2:**
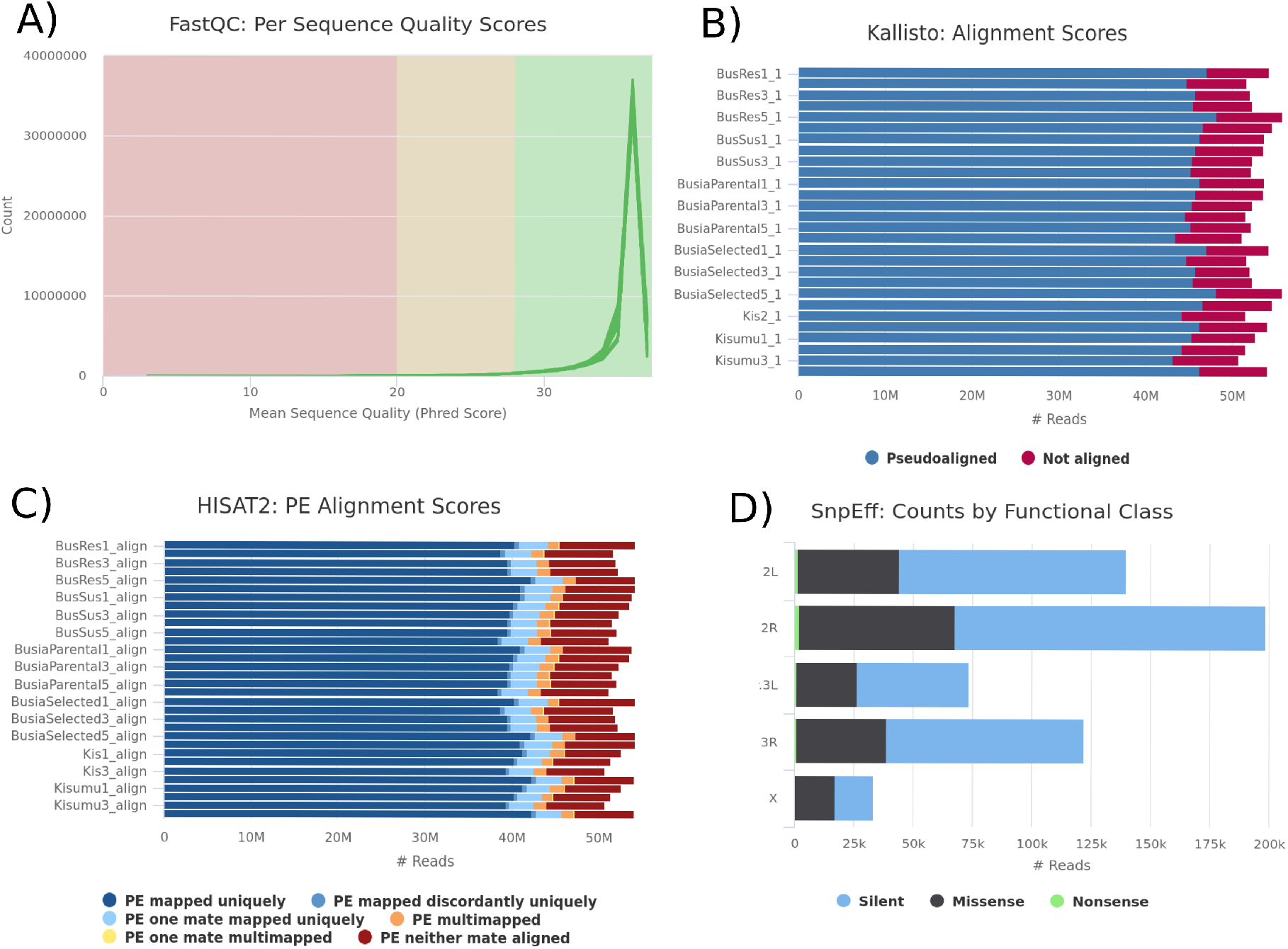
MultiQC captures quality control statistics from across the *RNA-Seq-Pop* workflow. a) per-base sequence content as calculated by FASTQC b) Total reads and number of successfully aligned reads to the reference transcriptome by Kallisto. c) The number of reads that were successfully mapped to the reference genome with HISAT2 d) The proportion of missense, synonymous and nonsense SNPs reported by snpEff.

We removed adapter sequences from the paired-end reads and aligned them to the *Anopheles gambiae*PEST reference transcriptome (AgamP4.12) (Figure 2b). 844.25 million reads were processed in total, with 727.84 million successfully aligned, giving an overall 85.58% alignment rate (+/− 0.206% standard error) across sixteen total replicates. The breakdown of reads counted per sample can be found in supplementary Figure 3.

As a further quality control step, and to uncover the overarching relationships of gene expression between samples, *RNA-Seq-Pop* performs a principal components analysis (Figure 3a), and a sample-to-sample correlation heatmap (Figure 3b) on the DESeq2 normalised count data. In both analyses, biological replicates of each treatment group clustered together, supporting robust replication in these samples.

**Figure 3:**
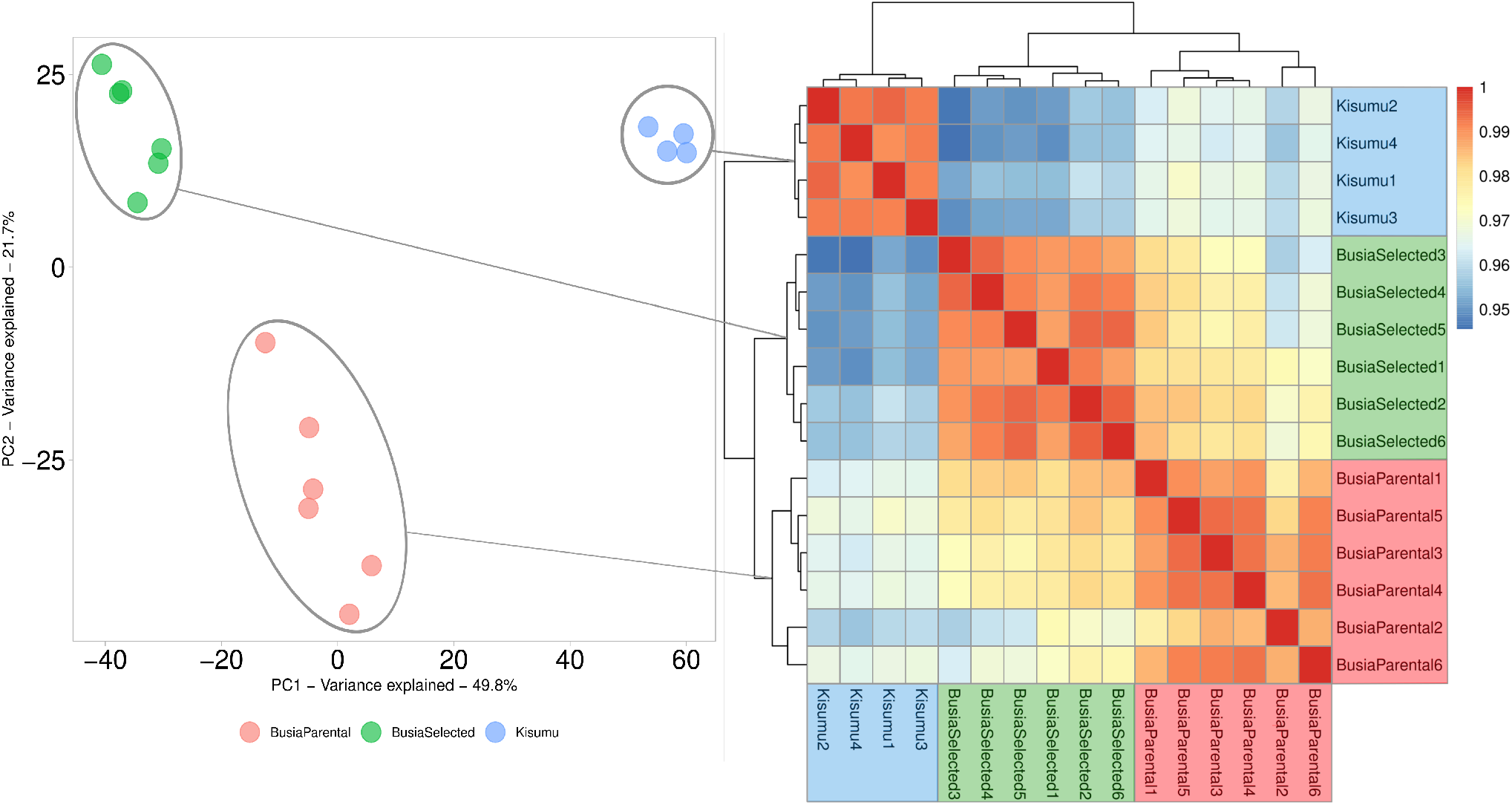
Exploratory sample clustering. a) Principal Components Analysis of the normalised read count data, showing clear separation between conditions b) A sample-to-sample Pearson’s correlation heatmap of normalised read counts assigned to each biological replicate, dendrograms show heirarchical clustering applied directly to Pearson’s correlations

#### Differential expression

We compared the selected Busia strain primarily to the parental strain, and also to the lab-susceptible Kisumu, which provides a cross-reference with earlier studies, as well as an extra level of filtering to identify candidate genes. Our DESeq2 differential expression analysis (Wald test) identified 5416 differentially expressed genes between Kisumu and the parental Busia line and 5657 between the parental Busia and selected Busia. The full table of differentially expressed genes in all comparisons can be found in the supplementary file S1, and volcano plots in supplementary figures 4a, b, c.

The high sequencing depth performed provides ample power to detect differences in expression. For example, a number of genes belonging to candidate detoxification families that are known to interact with insecticides were significantly differentially expressed, for example, 51 cytochrome P450s, 23 carboxylesterases and 20 ABC transporters. All three sensory appendage protein (*Sap)* genes in the *An. gambiae* genome were significantly overexpressed in the selected Busia strain compared to the parental Busia line. *Sap2* showed 10.7 fold overexpression (6.5-17.5 95% CIs), while *Sap1* exhibited 1.8-fold (1.36-2.44 95% CIs) and *Sap3* 2-fold (1.58–2.51 95% CIs) overexpression.

Using an option within *RNA-Seq-Pop* that compares expression trends across multiple comparisons, we identified a cluster of carboxylesterases which were overexpressed in Busia (G24) vs Kisumu and in Busia (G28) vs Busia (G24). In the latter comparison, *Coebe2c* showed a fold change of 1.69 (1.3-2.1 95% CIs), *Coebe3c* 3.05 (1.6-5.9 95% CIs) and *Coebe4c* 1.61 (1.2-2.2 95% CIs). We examined whether any selective sweeps were observed around these loci in the Ag1000g phase 1 data set and identified one in the *An. gambiae* Gabon population, although not in the Ugandan sample.

#### Variant calling

We enabled *RNA-Seq-Pop* to call genomic variants with *freebayes* and output data in VCF format. Across all chromosomes, and after filtering, *RNA-Seq-Pop* called 734,269 variants. Figure 5 shows a visual representation of genome composition in the *Anopheles gambiae PEST* reference genome, and the proportion of SNPs covered by each genomic feature in our genotype calls. The *An. gambiae* genome consists of 54%intergenic and 46% genic sequence (of which 14% are exonic, and 32% intronic). Given the nature of RNA-Seq, we expected to primarily find SNPs in coding regions of the genome, which are expressed. Indeed, of these 734,269 variants, we find 73% residing within exons, 11% in introns, and 16% in intergenic regions. The finding of 16% of SNPs in intergenic regions is likely to be explained by expression of non-coding RNAs, and the misannotation of transcripts – particularly 5’ and 3’ UTRs. The workflow automatically annotates the called variants with snpEff – across all exons, 16.4% of variants were annotated as non-synonymous, and 58.1% as synonymous.

**Figure 4:**
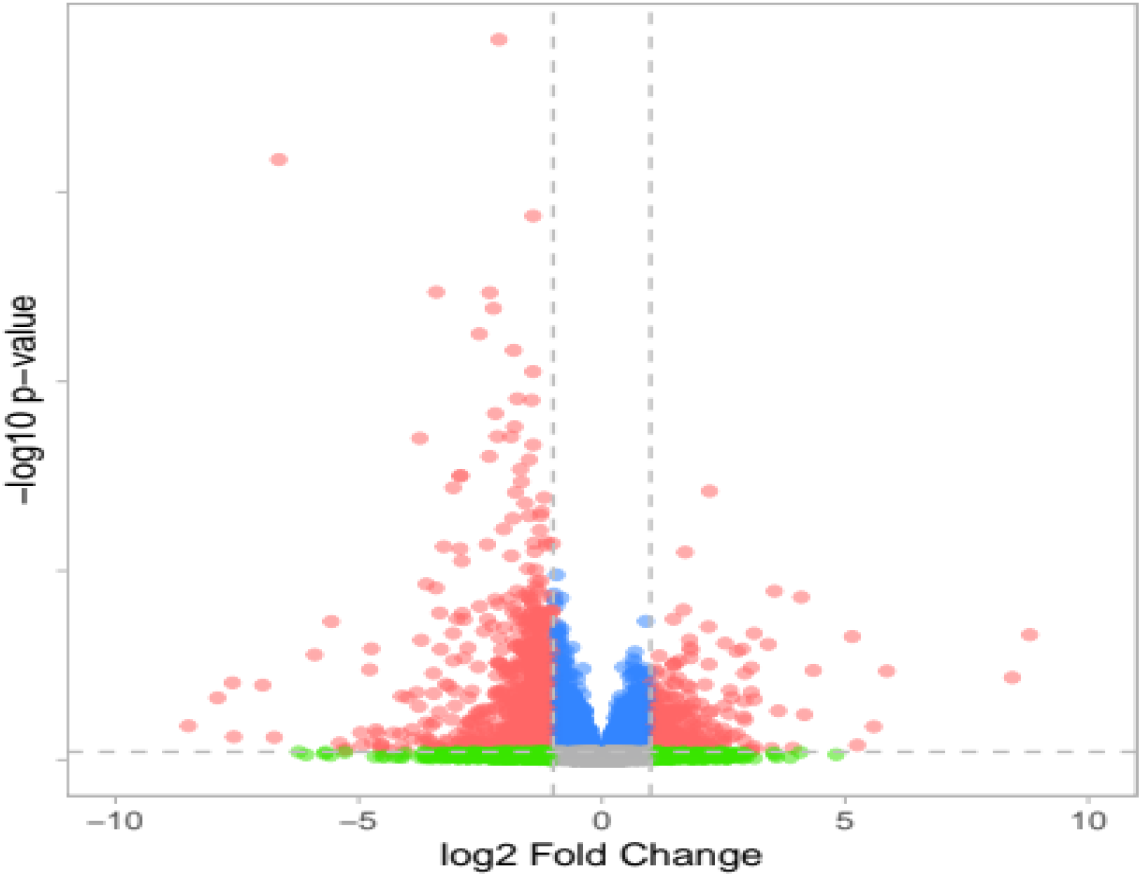
A volcano plot showing gene expression differences between Busia Parental and Busia Selected. -Log10 P-values are plotted against Log2 Fold Change. Red=genes with adjusted p-value < 0.05 and an absolute fold change > 2, green= adjusted pvalue > 0.05 and a fold change > 2, blue= adjusted pvalue > 0.05 and a fold change < 2. An outlier (AGAP012637) has been removed for visualisation purposes.

**Figure 5:**
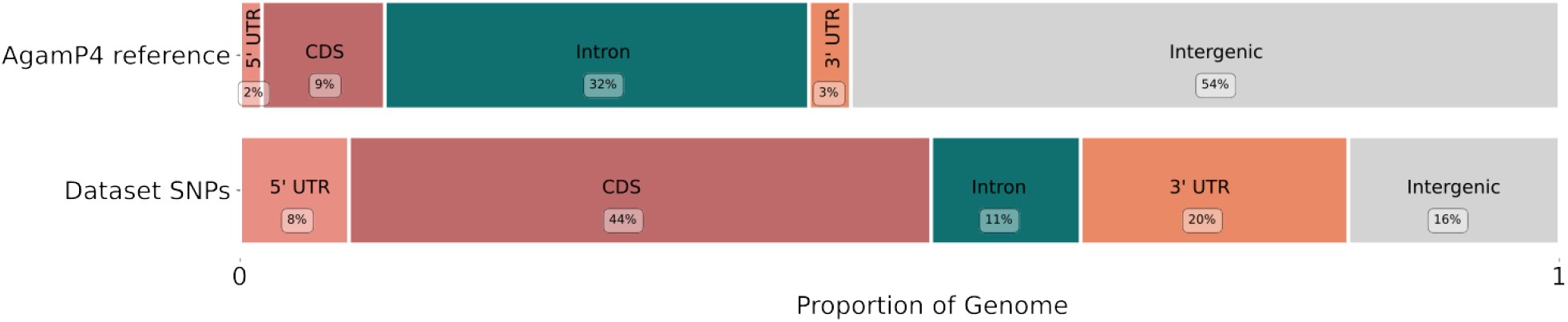
SNPs from RNA-Seq are enriched in transcribed regions. Illustration of the proportion of SNPs found within each genomic feature in the AgamP4 reference genome (Upper panel) and in the combined Busia and Kisumu *Anopheles gambiae* RNA-Seq dataset (Lower panel).

There was a positive correlation between read counts per gene, and the number of called SNPs per gene when controlling for gene size (GLM – coef=0.135, pval=2.2e^-36^, supplementary Table 5). A PCA based upon read count data, was not qualitatively different from the PCA on expression data (Figure 3 and supplementary Figure 6).

#### Genetic diversity

Table 1 shows genome-wide nucleotide diversity (π) and Watterson’s θ, averaged across 20kb nonoverlapping windows. To standardise sample size we down-sampled both Busia strains from six to four replicates. Both measures of genetic diversity were significantly lower in the Kisumu strain compared to the two Busia strains, as would be expected after a long history of laboratory colonisation. The selected Busia line also shows a reduction in genetic diversity compared to its founding strain, the parental Busia colony.

**Table 1:** Genetic Diversity. Average measures of genetic diversity, calculated in 20kb overlapping windows, across chromosomal arms. a) π, Nucleotide diversity b) Θ, Watterson’s theta

| | $\pi$ (95% CIs) | $\theta$ (95% CIs) |
| --- | --- | --- |
| Busia Parental | $1.04 \times 10^{-3}$ ($1.02 \times 10^{-3}$-$1.07 \times 10^{-3}$) | $7.4 \times 10^{-4}$ ($7.23 \times 10^{-4}$-$7.57 \times 10^{-4}$) |
| Busia Selected | $7.07 \times 10^{-4}$ ($6.87 \times 10^{-4}$-$7.27 \times 10^{-4}$) | $5.51 \times 10^{-4}$ ($5.37 \times 10^{-4}$-$5.65 \times 10^{-4}$) |
| Kisumu | $6.18 \times 10^{-4}$ ($6.0 \times 10^{-4}$-$6.35 \times 10^{-4}$) | $4.06 \times 10^{-4}$ ($3.95 \times 10^{-4}$-$4.18 \times 10^{-4}$) |

#### Known insecticide resistance variants of interest

If provided with a list of user-defined variants of interest, *RNA-Seq-Pop* will generate reports and plots of allele balance (the allele frequency found in the read alignments). For our variants of interest, we curated a selection of SNPs which have been associated with insecticide resistance in previous studies. Figure 6 shows allele frequencies of variants of interest across all samples. We show that over the four generations of selections, the frequency of the *Vgsc*-995S *kdr* allele increased from 25% (95% CIs: 21.5-29.8%) in G24 to fixation (100%) in the selected G28 Busia strain. In agreement with recent work from the Ag1000g project, we found no known secondary *kdr* mutations alongside the *Vgsc*-995S allele (Clarkson *et al.,* 2021).

**Figure 6:**
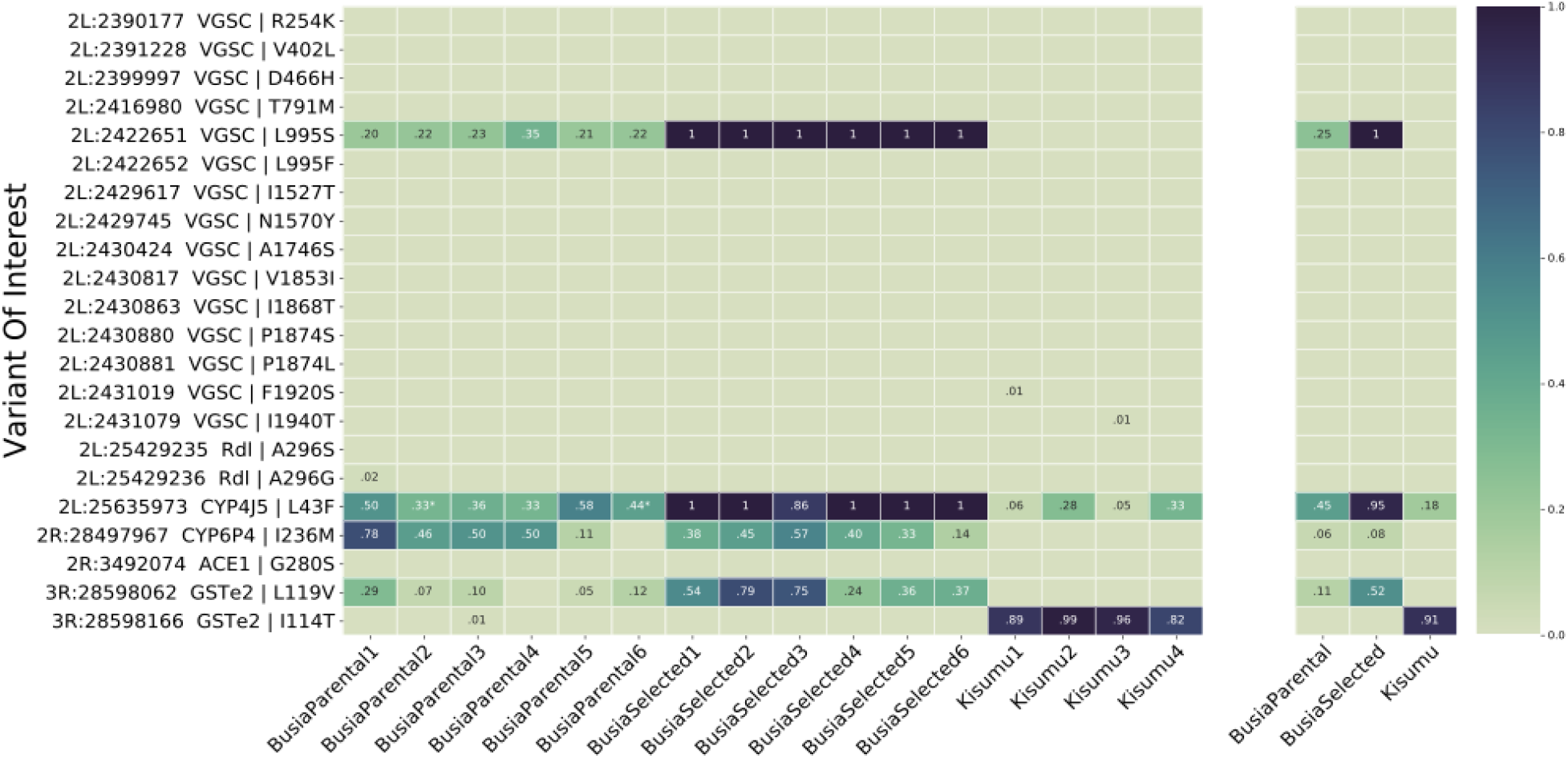
Variants of Interest. A heatmap showing allele frequencies of variants of interest found in read data in a) each sample and b) overall average allele frequency across strains. Blank cells indicate that the mutant allele was not detected despite reads across that genomic position.

In addition, the *Cyp4j5*-L43F mutation, previously associated with insecticide resistance in Uganda, showed a large increase in frequency after the selection regime, increasing from an average frequency of 43% (95% CIs: 32-54%) to 98% (95% CIs: 94-100%). The *Gste2-I114T* mutation, associated with DDT resistance, was absent in both Busia strains, however surprisingly, it was present at high frequency (92%) in the pyrethroid susceptible Kisumu reference strain. Another mutation, *Gste2*-L119V, increased in frequency from 11% (95% CIs: 9-13%) to 52% (95% CIs: 47-58%). The *Cyp6p4-I236M* mutation, linked to a swept haplotype in Uganda, was present in Busia samples, but there was no significant difference in frequency between the parental (39%, 95% CIs: 29-53%) and selected groups (38%, 95% CIs: 26-52%). In agreement with these differences in frequency of known insecticide-resistance variants, we find *F_st_* values in both the *Vgsc* and *Cyp4J5* genes in the top 5% percentile between the G24 parental Busia strain and the G28 selected Busia strain, but not in *Cyp6P4* (89^th^ percentile).

The *Ace-1-*G280S mutation was absent from all samples. A single allele of the *rdl*-A296G mutation was detected in the Parental Busia strain, however, this could have resulted from a base-calling error in the sequencing reads. Complete allele balance data for all variants of interest can be found in the supplementary file S2. We looked within the primary candidate gene from differential expression analysis, *Sap2,* for allele frequency changes, but no non-synonymous variants were present in the data.

#### Selection

The workflow permits calculation of *F_st_* and the population branch statistic (PBS) both in windows as genome-wide selection scans (GWSS) and within each gene. In the context of insecticide resistance, finding regions of high genetic differentiation between susceptible and resistant mosquito populations can allow us to identify loci or variants that contribute to the phenotype. We found high overall levels of *F_st_* between the parental Busia and the selected Busia, however, *F*_st_ on chromosomal arm 2L was especially elevated as compared to the other arms (supplementary Table 8). In the Busia data, the GWSS’s exhibit a large degree of noise, which may result from the inbred nature of the colonies used in this analysis. In other datasets from F1 *An. gambiae* (examples in supplementary figure 11), the genome-wide selection scans are able to capture signals at sites of known selective sweeps.

#### Chromosomal Inversions

We estimated the karyotype of the samples with *compkaryo* for the 2La and 2Rb chromosomal inversions, by extracting karyotype-tagging SNPs. We focus on these two inversions because both contain a large number of tagging SNPs, providing confidence in the overall calls. Figure 7 shows a diagram of the *An. gambiae* genome, with the location and average karyotype frequency per group. After the four generations of selections, the 2La inversion rose significantly in frequency from an average of 33% to 86% (Mann-Whitney U, Adjusted P-value = 0.014), where 0% means no 2La alleles across all tagSNP loci, and 100% means all 2La alleles across all tagSNP loci. The frequency of the 2Rb inversion was also significantly different between Kisumu and both Busia colonies (Mann-Whitney U, Adjusted P-values < 0.05). Supplementary figure 10 shows the per-replicate karyotype frequency.

**Figure 7:**
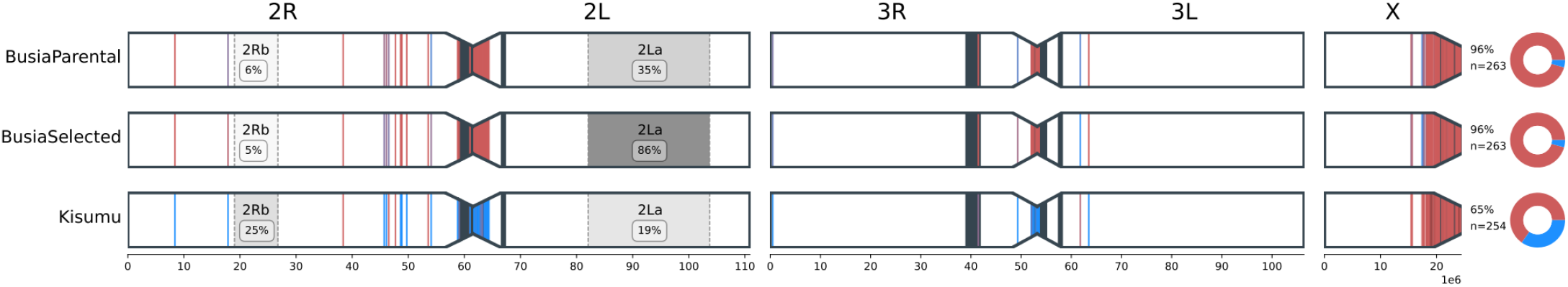
Ancestry and karyotyping. Left) A diagram of the mosquito chromosomal arms, including heterochromatin regions (black). Ancestry informative markers that are indicative of either *An. gambiae* (red) or *An. coluzzii* (blue) are displayed as vertical lines. The major inversions 2La and 2Rb are displayed, along with their respective average frequency amongst treatment groups, as called by the program compkaryo. Right) A donut chart of the proportion of ancestry informative markers that are indicative of either *An. gambiae* (red) or *An. coluzzii* (blue) ancestry for each sample. The overall proportion of gambiae alleles (%) and the number of called AIMs (n=) per group is labelled.

#### Ancestry

Ancestry informative markers are SNPs which show fixed (or almost fixed) differences between species. *RNA-Seq-Pop* can utilise sets of Ancestry Informative markers to investigate the proportion of ancestry for each chromosome assigned to either *An. gambiae, An. coluzzii or An. arabiensis.* Figure 7 shows the position of called AIM alleles that map to either *An. gambiae* or *An. coluzzii* across the genome. This shows that the Busia samples were primarily of *An. gambiae s.s* ancestry across all chromosomes, in concordance with the X chromosome-based SINE species ID assay (Santolomazza et al., 2008). However, the pattern was markedly different for the susceptible reference strain, Kisumu, which showed a large degree of putative *An. coluzzii* ancestry on the autosomes (supplementary table 9).

## Discussion

### *RNA-Seq-Pop* Implementation

*RNA-Seq-Pop* encompasses a complete workflow for RNA-Sequencing analysis, from quality control and read trimming, to transcript quantification and differential expression analysis (DEA). However, as well as conducting traditional differential expression analyses at both the gene and isoform level, *RNA-Seq-Pop*exploits useful, but often ignored sequence data.

*RNA-Seq-Pop* is designed for ease of use, requiring only a *sample metadata sheet* and a yaml format *configuration* file. A single command in the terminal will automatically install all dependencies and run the workflow, which is scaled by Snakemake to run on a personal computer, cluster or cloud environment. The workflow is applicable to any Illumina paired-end RNA-Sequencing data, and is flexible, allowing for variation in ploidy; including haploid, diploid, or pooled samples. We have written *RNA-Seq-Pop* in accordance with Snakemake best practices (Köster, 2022), and hope that it is an intuitive program, readily configured by the user to allow reproducible transcriptomic analyses. To increase accessibility *RNA-Seq-Pop* is written in python and R, the two most popular programming languages in the life sciences.

Decreasing sequencing costs have facilitated the proliferation of genomic surveillance in infectious disease research (Neafsey et al., 2021). The specific modules within *RNA-Seq-Pop*, which are readily adapted to other organisms, allow us to investigate novel variants that may be involved in our phenotype of interest (insecticide resistance), while simultaneously producing data on known resistance variants which can provide actionable information for malaria control programme personnel. For *An. gambiae s.l,* we provide a versioned variants of interest file in the GitHub repository, which we will update with additional resistance or disease transmission-related variants. As well as highlighting known variants, *RNA-Seq-Pop* can also perform genome-wide selections scans, using *Fst* (Bhatia et al., 2013) and the Population Branch statistic, PBS (Yi et al., 2010), highlighting known and novel regions of the genome that may be involved in the phenotype of interest.

For the major malaria vector, *An. gambiae s.l, RNA-Seq-Pop* can determine the karyotype frequency of chromosomal inversions, utilising the program *compKaryo* (Love et al., 2019). *An. gambiae s.l* has been shown to harbour a number of segregating chromosomal inversions, which have been associated with environmental heterogeneity, susceptibility to *Plasmodium* infection, and with insecticide resistance (Coluzzii et al., 1979, Riehle et al., 2017, Weetman et al., 2018). Typically, we can only detect these inversions through molecular PCR-based assays (of which many do not exist for the range of inversions karyotyped by *compKaryo*) or laborious and technically challenging cytologic experiments (Coluzzi et al., 2002, White et al., 2007), although recent approaches using tagging SNP panels appear promising (Love et al., 2020).

We can also illuminate the putative ancestry of our samples. This is of particular interest as the two recently-diverged sibling species *An. gambiae* and *An. coluzzii,* may often hybridise, and have undergone extensive introgression in the recent past (Fontaine et al., 2015; Vicente et al., 2017), allowing resistance alleles to cross from one species to another (Clarkson et al., 2014; Grau-Bové et al., 2020, 2021). Despite this, molecular assays typically target only a single marker on the X chromosome, ignoring the potential for admixture elsewhere in the genome (Caputo et al., 2021; Chabi et al., 2019; Santolamazza et al., 2008).

### Patterns of resistance in the Busia dataset

The differential expression analysis highlighted a multitude of detoxification genes overexpressed in the selected Busia line, including cytochrome P450s, carboxylesterases, chemosensory proteins, and ABC transporters, reflecting the polygenic nature of insecticide resistance. Many P450 genes were ≈2 fold overexpressed and it is not known whether this is due to constitutive differences between the strains, or induction by deltamethrin exposure in the G28 Busia strain. The *Sap2* gene in particular was highly overexpressed (10.7 fold after deltamethrin selections), and thus serves as a strong candidate for pyrethroid resistance outside of the West African *An. coluzzii* populations in which it was originally identified (Ingham et al., 2020).

The fixation of the *Vgsc*-995S *kdr* allele following selection is as predicted given its known association with pyrethroid resistance. Interestingly, the selected Busia strain shows a much stronger phenotype against permethrin than deltamethrin, which could partially be a result of this mutation. Earlier studies have shown a stronger protective effect of the *Vgsc*-995S allele against permethrin than deltamethrin (Lynd et al., 2010). In agreement with this shift in *Vgsc*-995S frequency, we find high *Fst* in the *Vgsc* between the parental and selected Busia colonies. The *Vgsc* is not differentially expressed between the parental Busia strain and the selected Busia, meaning this result would have been missed using differential expression analyses alone.

During deltamethrin selections, the 2La inversion rose in frequency dramatically, suggesting an association with deltamethrin resistance in Busia. Associations between the 2La inversion and insecticide resistance have been previously reported (Weetman et al., 2018). We also find a large shift in *Cyp4J5-L43F* mutation frequency, which lies within the 2La inversion and a very high *F_st_* in this gene (0.59). Interestingly, the gene is also differentially expressed, perhaps suggesting that the 2La haplotypic background results in over-transcription of the gene when compared to 2L+a haplotypes. It is not clear whether *Cyp4J5* is causative, or if there are other variants on the 2La haplotype(s) that are driving this shift in 2La. In agreement with this and the shift in *kdr*, we find high overall *F_st_* between the parental and selected Busia lines on the 2L chromosomal arm (supplementary Table 8).

Interestingly, *RNA-Seq-Pop* revealed that the Kisumu reference strain, exhibits a large proportion of putative *An. coluzzii* ancestry. The Kisumu reference strain was colonised from Kisumu, Kenya in 1975 (Williams et al., 2019) from an area where *An. coluzzii* has not been recorded. The most parsimonious explanation is that the colony has been contaminated through hybridization in the insectary during its long colonisation. The X chromosome is typically resistant to introgression, and consistent with a theory of a lab contamination event no *An. coluzzii* variants are found on the X chromosome. The X chromosome of Kisumu also has a particularly low estimate of Watterson’s Θ compared to the autosomes, which may reflect admixture present on the autosomes (supplementary table 7A). In addition, we also find that the Kisumu strain contains the *Gste2-114T* mutation at high frequency. In agreement with this finding, recent data shows intermittent resistance to DDT in this strain (Williams et al., 2019). We also observe some putative *An. coluzzii* alleles in the two Busia strains. Whilst we cannot rule out other explanations, this set of ancestry informative markers were derived from Mali, and therefore it is likely that some may not be truly informative of ancestry outside of this population.

Estimated population allele frequencies derived from RNA-Seq data may not accurately reflect DNA-based allele frequencies. Allele-specific expression is one cause of this, where two or more alleles in a diploid or polyploid may be expressed at different levels, causing an imbalance. Despite this, previous studies have shown a strong correlation between expressed and true allele frequencies, particularly at higher sequencing depth (Jehl et al., 2021; Lopez-Maestre et al., 2016; Oikkonen & Lise, 2017; Quinn et al., 2013). In this study, we performed RNASeq at a high sequencing depth, and therefore can have more confidence overall in our genotype calls and subsequent analyses. We recommend generally that for differential expression analyses, low coverage RNA-Sequencing is sufficient (10-25 million reads, or 5-13.5X coverage for *An. gambiae),*whereas for variant analyses, higher coverage is preferred (25-60 million reads, or 13.5-32.4X coverage for *An. gambiae*).

## Data accessibility

The workflow is hosted at https://github.com/sanjaynagi/rna-seq-pop. We welcome and encourage any feedback or contributions to *RNA-Seq-Pop.* The variant of interest file is versioned and is included in the GitHub repository. Raw sequence data is deposited at the ENA under BioProject PRJNA748581. The modified version of compKaryo is found here https://github.com/sanjaynagi/compkaryo.

## Author contributions

SCN, DW and MJD conceived and designed the study. SCN and AO performed all experiments and SCN analysed the data. SCN, DW and MJD drafted the manuscript, and all authors read and approved the final version.

