## Supplementary data for "*RNA-Seq-Pop:* Exploiting the sequence in RNA-Seq - a Snakemake workflow reveals patterns of insecticide resistance in the malaria vector *Anopheles gambiae*"

### Supplementary – Nagi *et al.*, 2022

1) Literature review – published studies of RNA-Sequencing in disease vectors

**Databases** – Web of Science
**Date** - 03/05/2022
**Search terms** –

1) (Anopheles OR Aedes OR Culex OR vector OR tsetse OR sandfly) AND (rna-seq OR rna-sequencing OR rna seq OR expression OR transcriptomics)
2) Mosquito AND (rna-seq OR rna-sequencing OR rna seq OR expression OR transcriptomics)

| Title | Citation | Taxon | Phenotype / purpose | Year | Sequence data utilised |
|---|---|---|---|---|---|
| The RNA-Seq approach to studying the expression of mosquito mitochondrial genes | (Neira-Oviedo et al., 2011) | *Aedes aegypti*, *Anopheles gambiae, and Anopheles quadrimaculatus* | Mitochondria | 2011 | |
| RNA-seq analyses of blood-induced changes in gene expression in the mosquito vector species, *Aedes aegypti* | (Bonizzoni et al., 2011) | *Aedes aegypti* | Bloodmeal | 2011 | |
| Comparative Transcriptome Analyses of Deltamethrin-Resistant and -Susceptible *Anopheles gambiae* Mosquitoes from Kenya by RNA-Seq | (Bonizzoni et al., 2012) | *Anopheles gambiae* | Insecticide resistance (pyrethroid) | 2012 | |
| RNA-Seq reveals early distinctions and late convergence of gene expression between diapause and quiescence in the Asian tiger mosquito, *Aedes albopictus* | (Poelchau et al., 2013) | *Aedes albopictus* | Diapause | 2013 | |
| The Developmental Transcriptome of the Mosquito *Aedes aegypti*, an Invasive Species and Major Arbovirus Vector | (Akbari et al., 2013) | *Aedes aegypti* | Life-stage | 2013 | |
| De novo transcriptome sequencing and sequence analysis of the malaria vector *Anopheles sinensis* | (Chen et al., 2014) | *Anopheles sinensis* | Annotation | 2014 | |
| Comparative analysis of response to selection with three insecticides in the dengue mosquito *Aedes aegypti* using mRNA sequencing | (David et al., 2014) | *Aedes aegypti* | Insecticide resistance (permethrin, imidacloprid, propoxur) | 2014 | Yes |
| Dual RNA-seq of Parasite and Host Reveals Gene Expression Dynamics during Filarial Worm–Mosquito Interactions | (Choi et al., 2014) | *Aedes aegypti* | Host-parasite | 2014 | |
| RNA-seq analyses of changes in the *Anopheles gambiae* transcriptome associated with resistance to pyrethroids in Kenya: identification of candidate-resistance genes and candidate-resistance SNPs | (Bonizzoni et al., 2015) | *Anopheles gambiae* | Insecticide resistance (pyrethroid) | 2015 | Yes |
| Comparative transcriptome analyses of deltamethrin-susceptible and -resistant *Culex pipiens pallens* by RNA-seq | (Lv et al., 2016) | *Culex pipiens* | Insecticide resistance (pyrethroid) | 2016 | |
| Single molecule RNA sequencing uncovers trans-splicing and improves annotations in *Anopheles stephensi* | (Jiang et al., 2017) | *Anopheles stephensi* | Annotation | 2017 | |
| Comparative Transcriptomics of Malaria Mosquito Testes: Function, Evolution, and Linkage | (Cassone et al., 2017) | *A. gambiae and A. merus* | Spermatogenesis | 2017 | |
| In the hunt for genomic markers of metabolic resistance to pyrethroids in the mosquito *Aedes aegypti*: An integrated next-generation sequencing approach | (Faucon et al., 2017) | *Aedes aegypti* | Insecticide resistance (pyrethroid) | 2017 | Yes |
| RNA-Seq Comparison of Larval and Adult Malpighian Tubules of the Yellow Fever Mosquito *Aedes aegypti* Reveals Life Stage-Specific Changes in Renal Function | (Li et al., 2017) | *Aedes aegypti* | Life-stage | 2017 | |
| The choreography of the chemical defensome response to insecticide stress: insights into the *Anopheles stephensi* transcriptome using RNA-Seq | (De Marco et al., 2017) | *Anopheles stephensi* | Insecticide resistance | 2017 | |
| Blood-induced differential gene expression in *Anopheles dirus* evaluated using RNA sequencing | (Mongkol et al., 2018) | *Anopheles dirus* | Bloodmeal | 2018 | |
| High-resolution transcriptional profiling of *Anopheles gambiae* spermatogenesis reveals mechanisms of sex chromosome regulation | (Taxiarchi et al., 2019) | *Anopheles gambiae* | Spermatogenesis | 2019 | |
| Transcriptome Sequencing and Analysis of Changes Associated with Insecticide Resistance in the Dengue Mosquito (*Aedes aegypti*) in Vietnam | (Lien et al., 2019) | *Aedes aegypti* | Insecticide resistance | 2019 | |
| Genome-wide gene expression profiling reveals that cuticle alterations and P450 detoxification are associated with deltamethrin and DDT resistance in *Anopheles arabiensis* populations from Ethiopia | (Simma et al., 2019) | *Anopheles arabiensis* | Insecticide resistance (pyrethroid) | 2019 | |
| UDP-glycosyltransferase genes and their association and mutations associated with | (Zhou et al., 2019) | *Anopheles sinensis* | Insecticide resistance | 2019 | |

| | | | | | |
|---|---|---|---|---|---|
| pyrethroid resistance in Anopheles sinensis (Diptera: Culicidae) | | | (pyrethroid) | | |
| RNASeq Analysis of *Aedes albopictus* Mosquito Midguts after Chikungunya Virus Infection | (Vedururu et al., 2019) | *Aedes albopictus* | Host-parasite | 2019 | |
| Contrasting patterns of gene expression indicate differing pyrethroid resistance mechanisms across the range of the New World malaria vector *Anopheles albimanus* | (Mackenzie-Impoinvil et al., 2019) | *Anopheles albimanus* | Insecticide resistance (pyrethroid) | 2019 | |
| Transcriptome analysis of *Anopheles dirus* and *Plasmodium vivax* at ookinete and oocyst stages | (Boonkaew et al., 2020) | *Anopheles dirus* | Plasmodium infection | 2020 | |
| Transcript Assembly and Quantification by RNA-Seq Reveals Significant Differences in Gene Expression and Genetic Variants in Mosquitoes of the *Culex pipiens* (Diptera: Culicidae) Complex | (Kang et al., 2021) | *Culex pipiens* | Insecticide resistance (pyrethroid) | 2021 | Yes |
| Integration of whole genome sequencing and transcriptomics reveals a complex picture of the reestablishment of insecticide resistance in the major malaria vector *Anopheles coluzzii* | (Ingham et al., 2021) | *Anopheles gambiae* | Insecticide resistance (pyrethroid) | 2021 | |
| Transcriptome comparison of dengue-susceptible and -resistant field derived strains of Colombian *Aedes aegypti* using RNA-sequencing | (Coatsworth et al., 2021) | *Aedes aegypti* | Vector competence | 2021 | |
| Transcriptomic and proteomic analysis of pyrethroid resistance in the CKR strain of *Aedes aegypti* | (Sun et al., 2021) | *Aedes aegypti* | Insecticide resistance (pyrethroid) | 2021 | |
| RNA-Seq analysis of blood meal induced gene-expression changes in *Aedes aegypti* ovaries | (Nag et al., 2021) | *Aedes aegypti* | Bloodmeal | 2021 | |
| Sympatric Populations of the *Anopheles gambiae* Complex in Southwest Burkina Faso Evolve Multiple Diverse Resistance Mechanisms in Response to Intense Selection Pressure with Pyrethroids | (Williams et al., 2022) | *Anopheles gambiae* | Insecticide resistance (pyrethroid) | 2022 | |
| RNAseq-based gene expression profiling of the *Anopheles funestus* pyrethroid-resistant strain FUMOZ highlights the predominant role of the duplicated CYP6P9a/b cytochrome P450s | (Wondji et al., 2022) | *Anopheles funestus* | Insecticide resistance (pyrethroid) | 2022 | |
| Transcriptome profiling reveals sex-specific gene expressions in pupal and adult stages of the mosquito *Culex pipiens* | (Martynova et al., 2022) | *Culex pipiens* | Life-stage | 2022 | |
| A whole transcriptomic approach provides novel insights into the molecular basis of organophosphate and pyrethroid resistance in *Anopheles arabiensis* from Ethiopia | (Messenger et al., 2021) | *Anopheles arabiensis* | Insecticide resistance | 2022 | Yes |

#### 2) Colony selection regime

Blood fed *Anopheles gambiae s.s* were collected from Busia, Uganda, in November 2018, using a prokopack aspirator. Approximately 200 mosquitoes were collected from 12 different homes in the village of South-Bugwere. Eggs were transported to the Liverpool School of Tropical Medicine and mosquitoes were reared in the insectaries at approximately 75% RH and 27°C, with a 12:12 hour light:dark photoperiod.

Between generations 10 and 24, due to the COVID-19 pandemic, Busia mosquitoes were not selected against deltamethrin to maintain their insecticide resistance status. As a result, the colony had lost resistance by G24, displaying 100% mortality to a 1 hour 0.05% deltamethrin (1X) WHO paper exposure, and 92.6% mortality to permethrin 0.75% (1X). Mosquitoes were then selected on 1X deltamethrin WHO papers for four consecutive generations (G24-G27), initially of 15 minute exposures, and subsequently for one hour.

##### A) Profiling

| Generation | Insecticide | Dead | Total | Mortality (%) |
|---|---|---|---|---|
| G24 | Deltamethrin | 99 | 99 | 100.0 |
| G24 | Permethrin | 88 | 95 | 92.6 |
| G25 | Deltamethrin | 47 | 54 | 87.0 |
| G25 | Permethrin | 15 | 47 | 31.9 |
| G26 | Permethrin | 9 | 30 | 30.0 |
| G27 | Deltamethrin | 164 | 217 | 75.6 |
| G28 | Deltamethrin | 145 | 208 | 69.7 |
| G28 | Permethrin | 23 | 106 | 21.7 |

##### B) Selections

1X deltamethrin (0.05%)

| Generation | Exposure. (mins) | Dead | Total | Mortality (%) |
|---|---|---|---|---|
| G24 | 15 | 1111 | 1205 | 92.2 |
| G25 | 15 | 348 | 456 | 76.3 |
| G26 | 60 | 265 | 327 | 81.0 |
| G27 | 60 | 164 | 217 | 75.6 |

**3)**

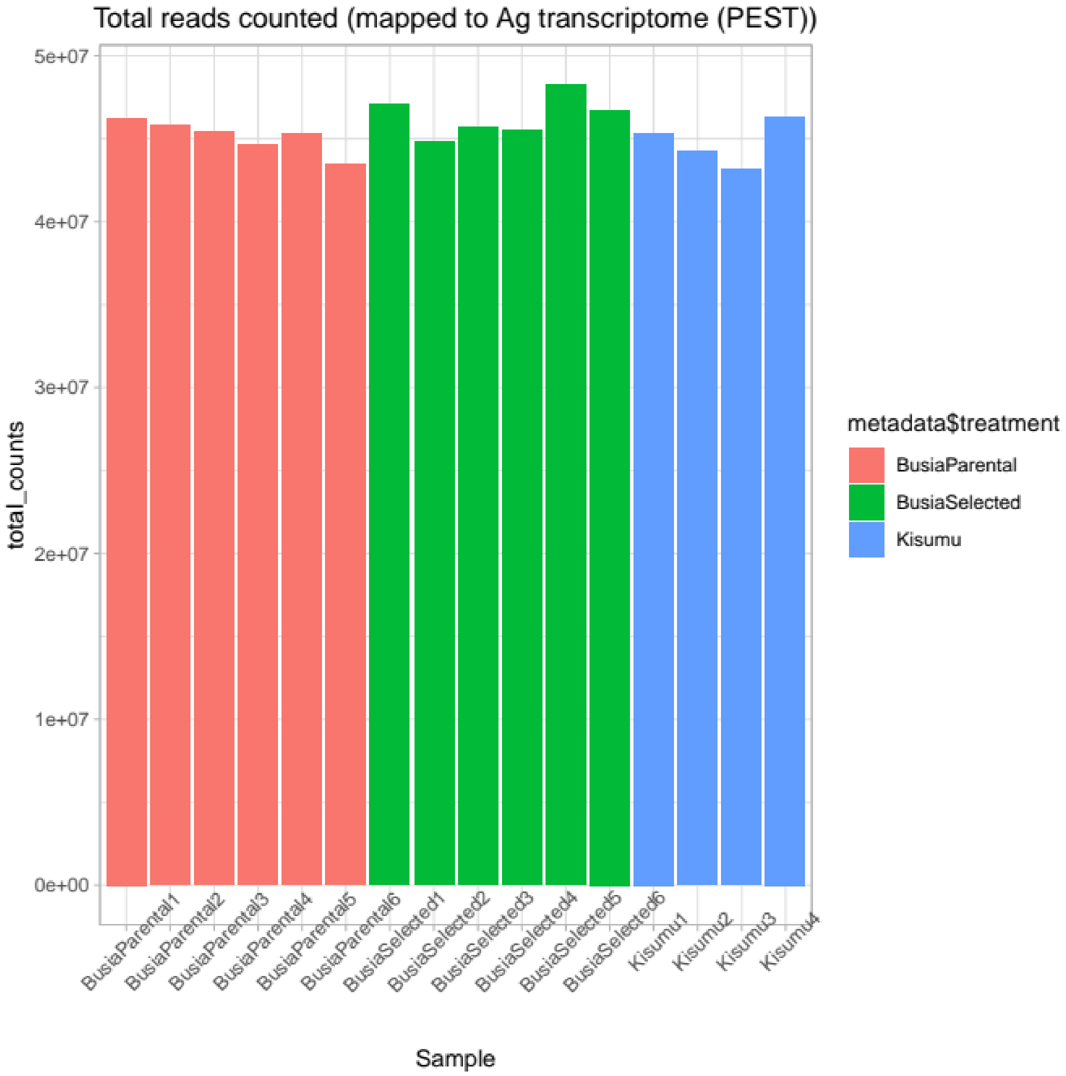
Total reads counted (mapped to Ag transcriptome (PEST))
total_counts
5e+07
4e+07
3e+07
2e+07
1e+07
0e+00
BusiaParental1
BusiaParental2
BusiaParental3
BusiaParental4
BusiaParental5
BusiaParental6
BusiaSelected1
BusiaSelected2
BusiaSelected3
BusiaSelected4
BusiaSelected5
BusiaSelected6
Kisumu1
Kisumu2
Kisumu3
Kisumu4
Sample
metadata$treatment
BusiaParental
BusiaSelected
Kisumu

### 4) Volcano plots

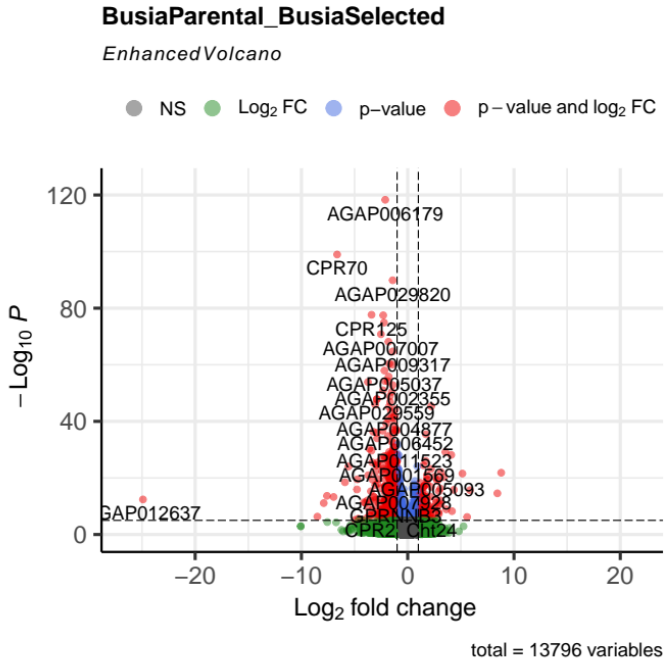


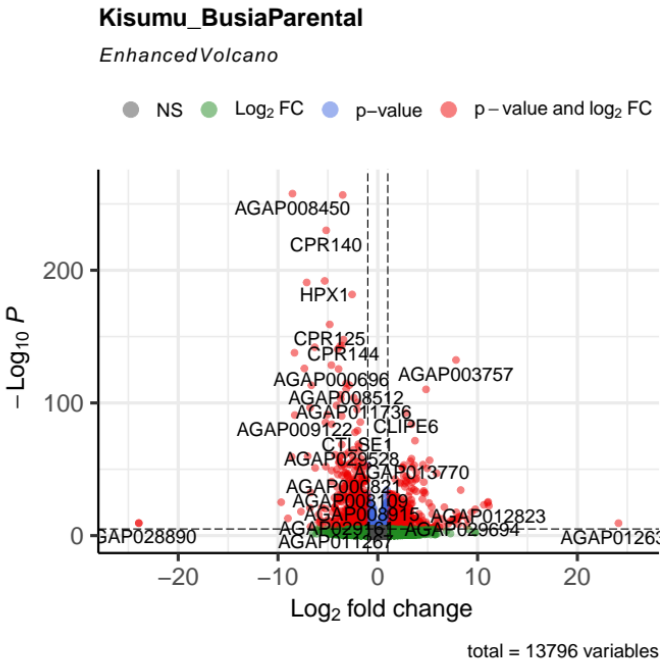


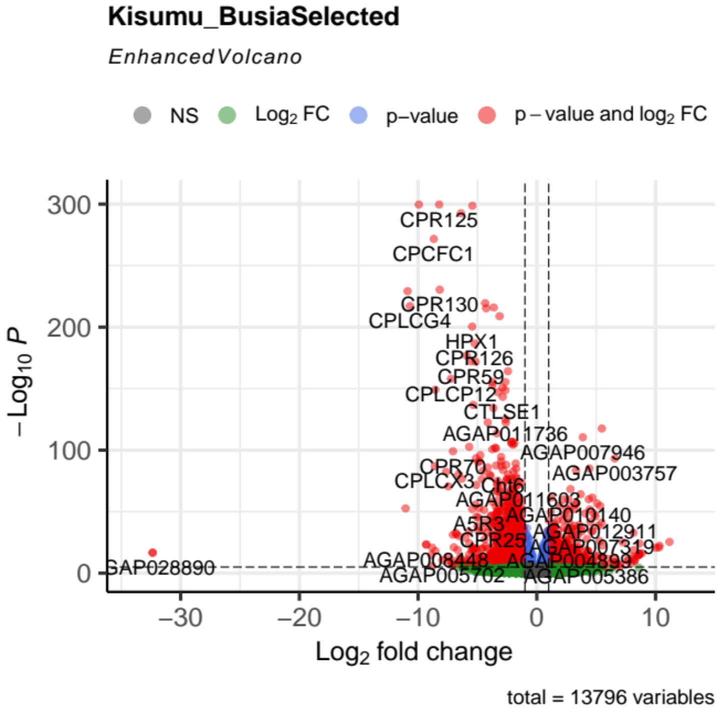

**5) OLS Regression of log2 number of SNPs per gene, total read counts per gene, and gene size in (bp).**

OLS Regression Results

| | | | |
|---|---|---|---|
| Dep. Variable: | nSNPs | R-squared: | 0.431 |
| Model: | OLS | Adj. R-squared: | 0.430 |
| Method: | Least Squares | F-statistic: | 449.4 |
| Date: | Fri, 21 Jan 2022 | Prob (F-statistic): | 5.48e-146 |
| Time: | 13:11:34 | Log-Likelihood: | -1734.4 |
| No. Observations: | 1189 | AIC: | 3475. |
| Df Residuals: | 1186 | BIC: | 3490. |
| Df Model: | 2 | | |
| Covariance Type: | nonrobust | | |

| | coef | std err | t | P>\|t\| | [0.025 | 0.975] |
|---|---|---|---|---|---|---|
| const | -2.4515 | 0.257 | -9.529 | 0.000 | -2.956 | -1.947 |
| Readcounts | 0.1462 | 0.011 | 13.698 | 0.000 | 0.125 | 0.167 |
| GeneSize | 0.4661 | 0.017 | 26.811 | 0.000 | 0.432 | 0.500 |

| | | | |
|---|---|---|---|
| Omnibus: | 87.964 | Durbin-Watson: | 1.592 |
| Prob(Omnibus): | 0.000 | Jarque-Bera (JB): | 106.486 |
| Skew: | -0.697 | Prob(JB): | 7.53e-24 |
| Kurtosis: | 3.457 | Cond. No. | 160. |

Notes:
[1] Standard Errors assume that the covariance matrix of the errors is correctly specified.

### 6) PCA

PCA X Ag_Busia

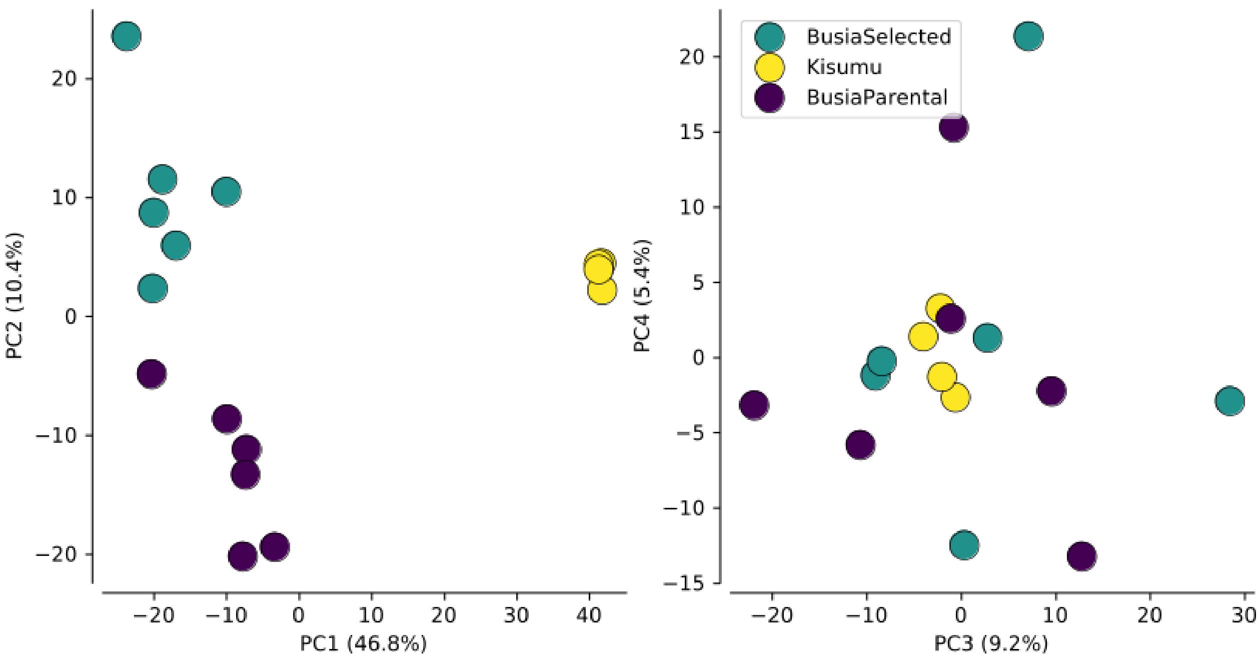


PCA 3R Ag_Busia

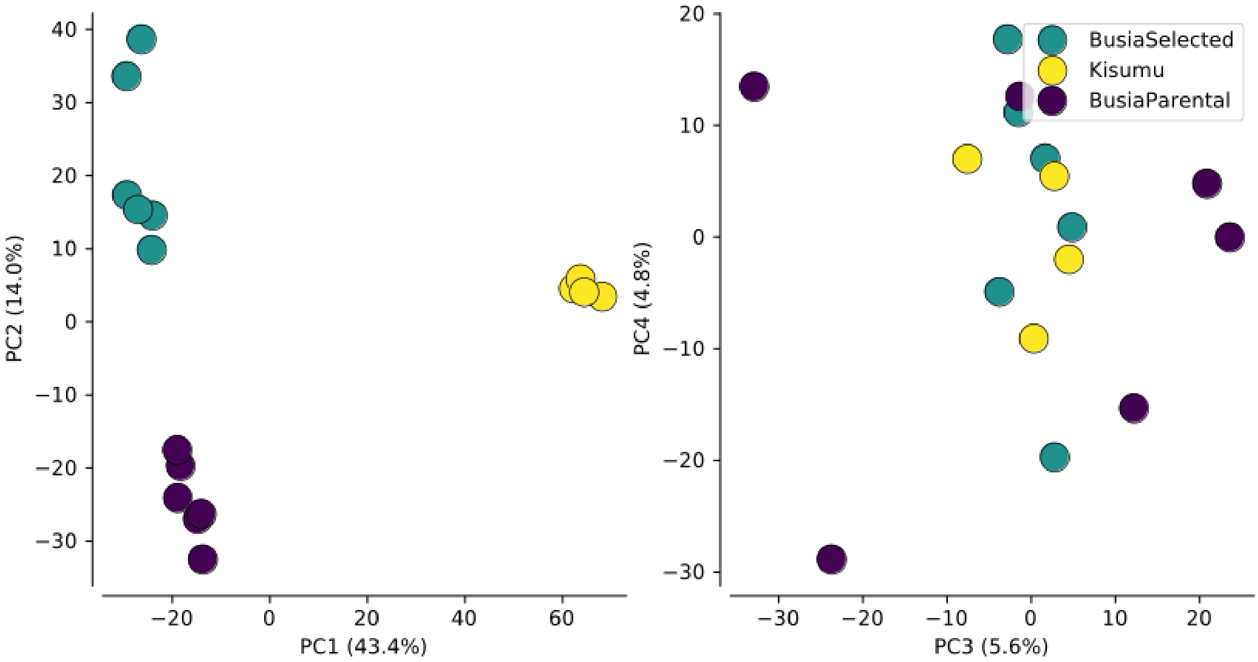

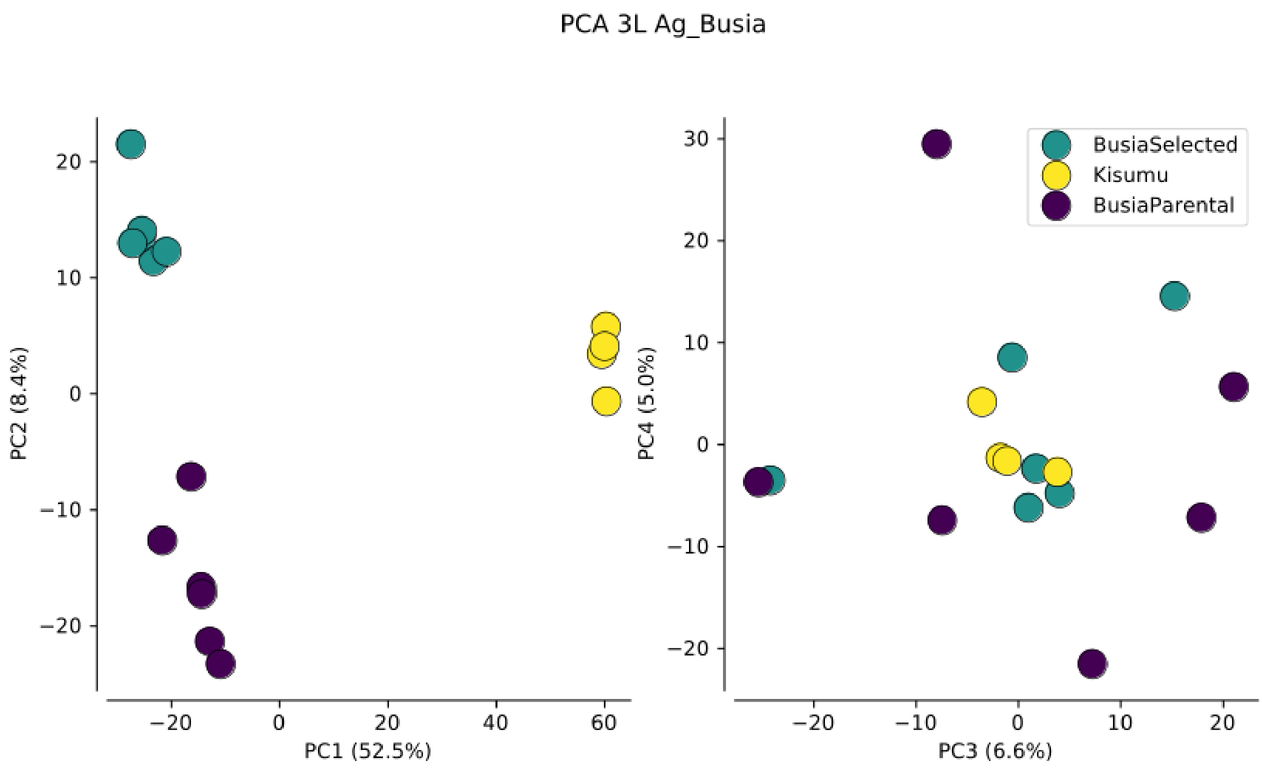
PCA 3L Ag_Busia
PC2 (8.4%)
PC1 (52.5%)
PC4 (5.0%)
PC3 (6.6%)
BusiaSelected
Kisumu
BusiaParental

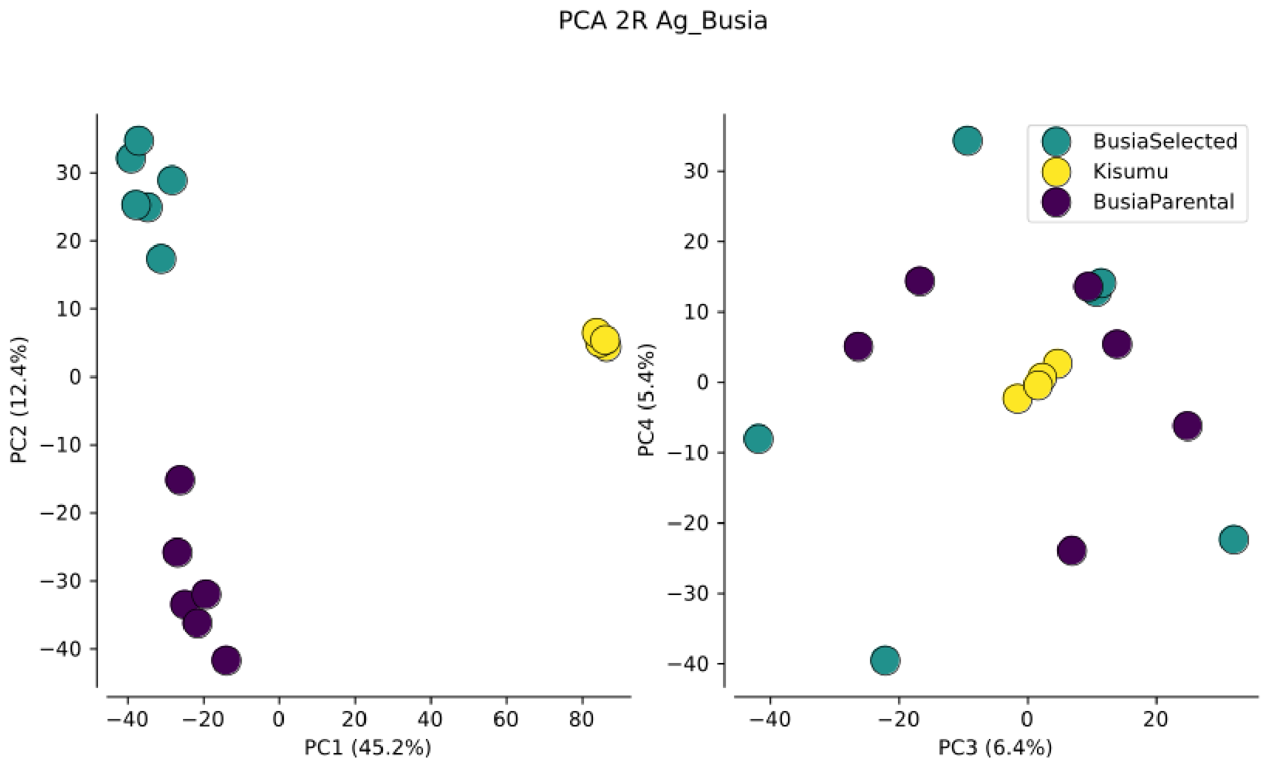
PCA 2R Ag_Busia
PC2 (12.4%)
PC1 (45.2%)
PC4 (5.4%)
PC3 (6.4%)
BusiaSelected
Kisumu
BusiaParental

PCA 2L Ag_Busia

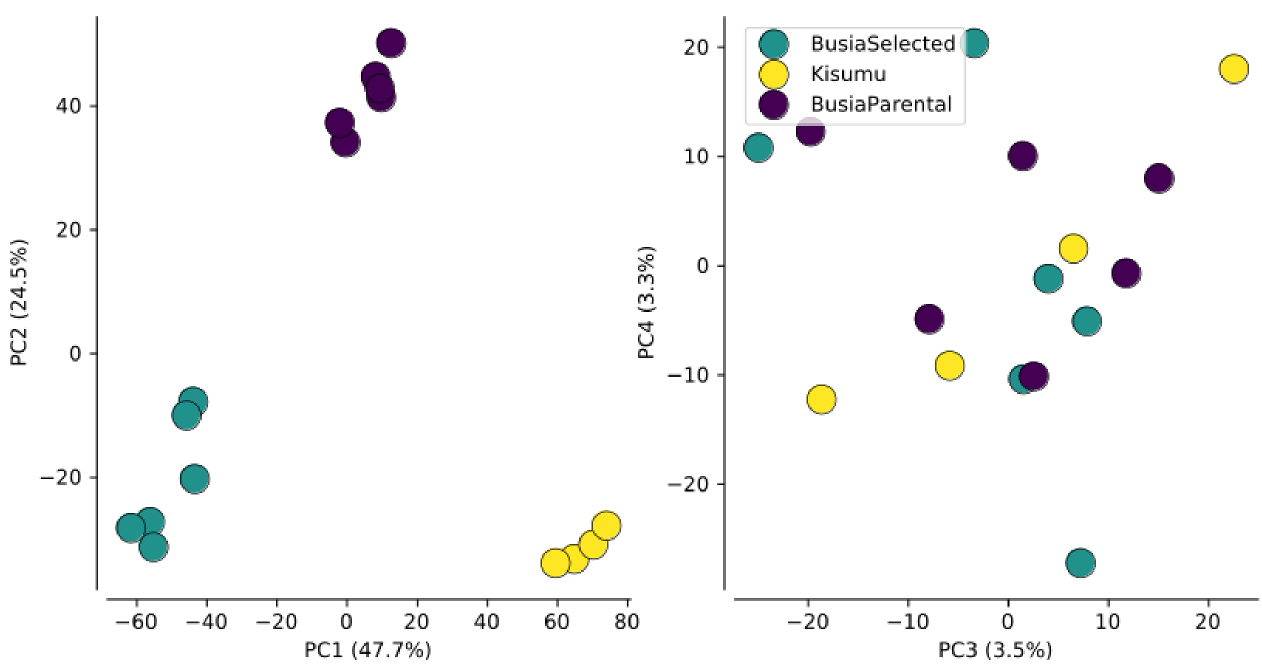

#### 7) Genetic diversity

**A)** Wattersons Theta

| Contig | Busia Selected | Busia Parental | Kisumu |
|---|---|---|---|
| **2L** | 0.00057 | 0.00068 | 0.00056 |
| **2R** | 0.00066 | 0.00083 | 0.00044 |
| **3L** | 0.00034 | 0.00059 | 0.00036 |
| **3R** | 0.00053 | 0.00067 | 0.00043 |
| **X** | 0.00068 | 0.00075 | 0.00025 |

**B)** Nucleotide Diversity

| Contig | Busia Selected | Busia Parental | Kisumu |
|---|---|---|---|
| **2L** | 0.00053 | 0.00102 | 0.00084 |
| **2R** | 0.00097 | 0.00116 | 0.00065 |
| **3L** | 0.00038 | 0.00069 | 0.00048 |
| **3R** | 0.00075 | 0.00092 | 0.00064 |
| **X** | 0.00097 | 0.00123 | 0.00034 |

8) Hudson Fst Per chromosome (Busia Parental v Busia selected)

| Contig | Fst |
| --- | --- |
| **2L** | 0.431 |
| **2R** | 0.109 |
| **3R** | 0.093 |
| **3L** | 0.11 |
| **X** | 0.106 |

#### 9) Proportion of ancestry per chromosome

| Strain | Contig | AIM fraction gambiae | AIM fraction coluzzii | n_aims |
|---|---|---|---|---|
| Busia Selected | 2L | 0.981 | 0.000 | 53 |
| Busia Selected | 2R | 0.796 | 0.093 | 18 |
| Busia Selected | 3L | 0.871 | 0.062 | 15 |
| Busia Selected | 3R | 0.836 | 0.164 | 16 |
| Busia Selected | X | 0.946 | 0.035 | 161 |
| Busia Parental | 2L | 0.981 | 0.000 | 53 |
| Busia Parental | 2R | 0.778 | 0.111 | 18 |
| Busia Parental | 3L | 0.887 | 0.047 | 15 |
| Busia Parental | 3R | 0.836 | 0.164 | 16 |
| Busia Parental | X | 0.945 | 0.036 | 161 |
| Kisumu | 2L | 0.105 | 0.875 | 51 |
| Kisumu | 2R | 0.185 | 0.703 | 18 |
| Kisumu | 3L | 0.057 | 0.877 | 15 |
| Kisumu | 3R | 0.170 | 0.830 | 16 |
| Kisumu | X | 0.966 | 0.014 | 154 |

#### 10) Karyotype frequencies calculated by compKaryo.

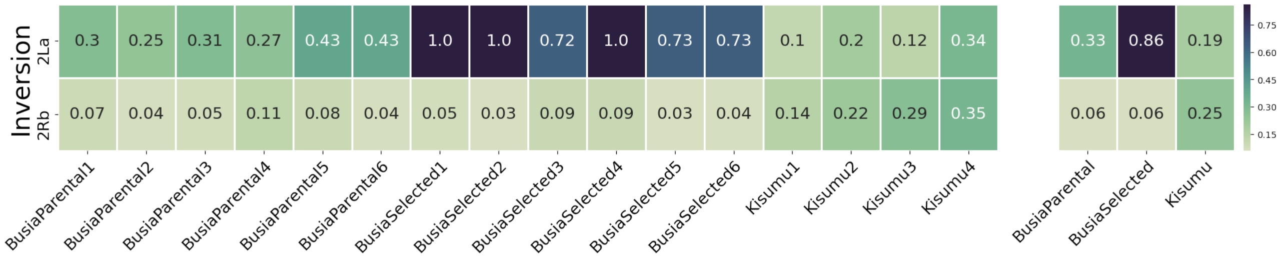


*Supplementary Figure 10: Karyotype frequencies. The frequency of the 2La and 2Rb karyotypes in a) each biological replicate and b) averaged across experimental conditions*

#### 11) GWSS – Fst

**Busia – chromosomal arm 2L**

Elevated Fst at the VGSC (beginning of chromosome)

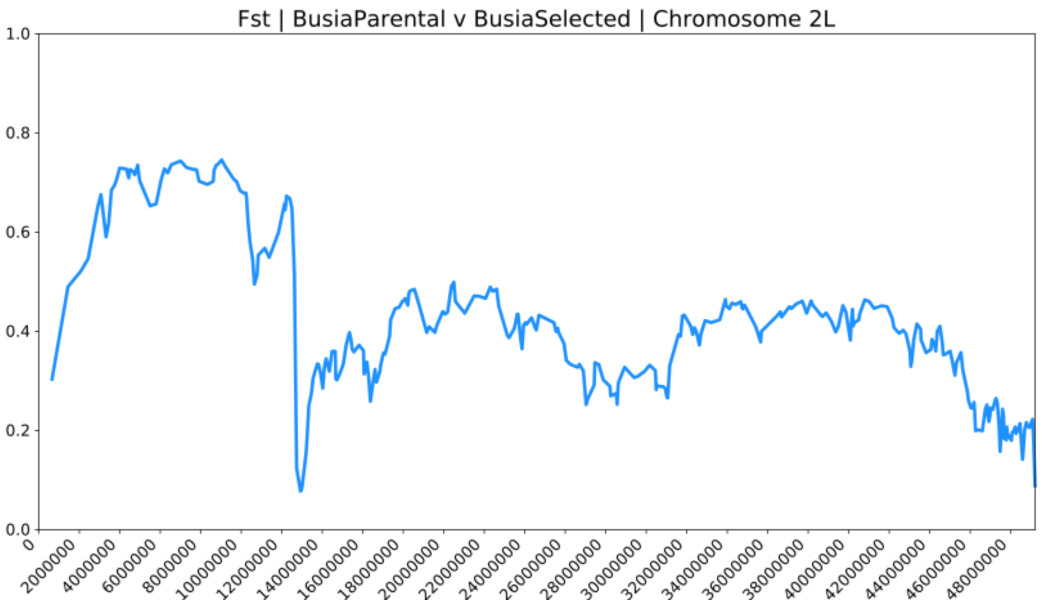


**Bouake F1 Population branch statistic - 2R**

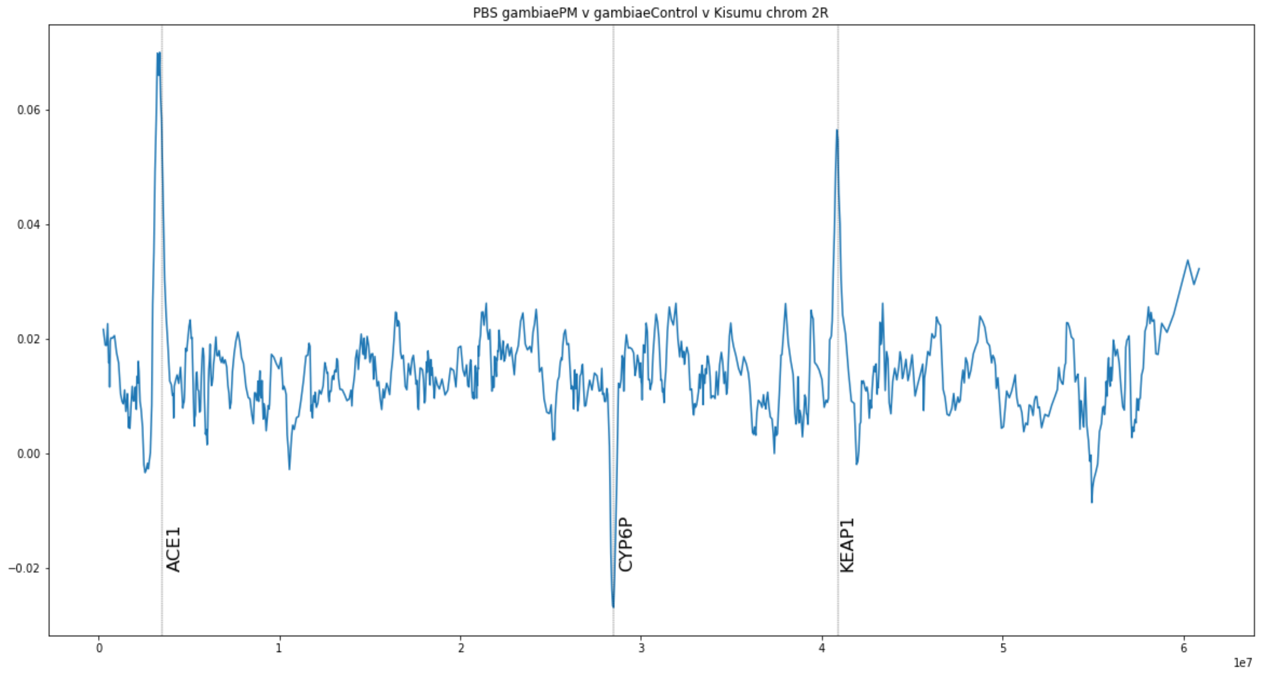
